# Nonameric structures of the cytoplasmic domain of FlhA and SctV in the context of the full-length protein

**DOI:** 10.1101/2020.10.12.336966

**Authors:** Lucas Kuhlen, Steven Johnson, Jerry Cao, Justin C. Deme, Susan M. Lea

## Abstract

Type three secretion is the mechanism of protein secretion found in bacterial flagella and injectisomes. At its centre is the export apparatus (EA), a complex of five membrane proteins through which secretion substrates pass the inner membrane. While the complex formed by four of the EA proteins has been well characterised structurally, little is known about the structure of the membrane domain of the largest subunit, FlhA in flagella, SctV in injectisomes. Furthermore, the biologically relevant nonameric assembly of FlhA/SctV has been infrequently observed and differences in conformation of the cytoplasmic portion of FlhA/SctV between open and closed states have been suggested to reflect secretion system specific differences. FlhA has been shown to bind to chaperone-substrate complexes in an open state, but in previous assembled ring structures, SctV is in a closed state. Here, we identify FlhA and SctV homologues that can be recombinantly produced in the oligomeric state and study them using cryo-electron microscopy. The structures of the cytoplasmic domains from both FlhA and SctV are in the open state and we observe a conserved interaction between a short stretch of residues at the N-terminus of the cytoplasmic domain, known as FlhA_L_/SctV_L_, with a groove on the adjacent protomer’s cytoplasmic domain, which stabilises the nonameric ring assembly.

## Introduction

Type III secretion systems (T3SS) are bacterial macromolecular machines that facilitate transport of protein substrates across the bacterial cell envelope (1, 2). The secretion machinery is conserved across at least two biological systems, the bacterial flagellum, which drives motility, and the virulence associated T3SS, which is an essential virulence factor for many bacterial pathogens due to the secretion of effector proteins directly into host cells (3). Most building blocks of the virulence associated T3SS or injectisome have a homologue in flagella and one of the most highly conserved parts of the system is a set of transmembrane proteins known collectively as the export apparatus (EA) (4).

The EA is made up of a core complex formed by the membrane proteins FliPQR (flagellar T3SS) or SctRST (virulence associated T3SS) which form a channel through which the unfolded protein substrates pass (5, 6). The regulatory subunit FlhB/SctU wraps around this core (7) and the inner rod and needle assemble onto its periplasmic face (8, 9). Surprisingly, following assembly into the full T3SS, the core complex is not in the plane of the inner membrane. Instead, it is held in the plane of the periplasm by interactions with the surrounding basal body and its hydrophobic surface is covered by another membrane protein complex situated directly underneath it in the inner membrane. This protein complex is believed to be formed by the transmembrane domain of the fifth EA component, FlhA or SctV (6, 10). The FlhA/SctV cytoplasmic domain is known to form a nonameric ring underneath FliPQR-FlhB/SctRSTU (11), but the full-length FlhA/SctV complex remains poorly characterised. The entire EA is housed within the basal body of the T3SS and is thought serve as a nucleus for its assembly (12, 13).

FlhA/SctV has long been known to be the major component of the EA, but its stoichiometry was controversial for a long time (14, 15) and may be dynamic *in vivo* (14, 16). Fluorescence based measurements showed that FlhA can form a large complex, but its exact stoichiometry couldn’t be measured with high precision (14, 16). The crystallisation of the cytoplasmic domain of SctV from *Shigella flexneri*, *Sf*-SctV, as a nonameric ring matching the dimensions of a toroidal density observed by tomography, established the stoichiometry of FlhA/SctV as nonameric (11). This was later confirmed by high resolution AFM, as the FlhA cytoplasmic domain also formed a nonameric ring on mica (17) whose stability depended on the linker between cytoplasmic and membrane domains, FlhA_L_/SctV_L_. Variability has been observed in the conformation of the cytoplasmic domains, with structural transitions proposed to be linked to the secretion process. Open and closed states have been described, with the open state being competent for chaperone-substrate complex binding (18), while the closed state has been proposed to be responsible for binding early substrates (19).

The transmembrane domain of FlhA/SctV is less well understood than the cytoplasmic domain. It is thought to contain approximately 8 transmembrane helices and conduct protons in order to use the pmf to power type III secretion (22). Consistent with this function, the transmembrane domain contains a number of conserved charged residues essential for function (23). Structural studies of the membrane domain have been hampered by the difficulties in producing the full-length membrane protein in the assembled state. In addition to the predicted transmembrane helices the membrane domain also contains a short stretch of highly conserved soluble residues known as the FHIPEP domain (24). It is known to be crucial for protein secretion, but its precise role in the secretion process remains to be elucidated (23, 25).

We produced assembled rings of full-length FlhA/SctV from both flagellar and virulence associated systems and studied them using cryo-EM. Here, we present the structures of the nonameric cytoplasmic domain of both FlhA and SctV. The structures demonstrate the conserved interaction between FlhA_L_/SctV_L_ and the neighbouring subunit and confirm the conserved stoichiometry of the complex in the context of the full length protein.

## Results

We screened a number of homologues of FlhA/SctV from many species for the ability to assemble in the absence of other export apparatus subunits in the *E. coli* membrane under overexpression conditions by employing superfolder GFP fusions (26) and assaying fluorescent spot formation in the cell envelope after overnight growth, suggesting successful membrane protein complex formation ^(14, 16, 27)^. We excluded FlhA/SctV homologues that only formed large spots at the cell poles, suggestive of protein aggregation (28). Most FlhA/SctV homologues formed distinct spots consistent with membrane localisation after overnight expression in *E. coli* BL21 cells, and the cells were observed to be highly elongated, consistent with disruption of the membrane machinery (Fig 1a). Out of the identified proteins we chose FlhA from the lateral flagella of Vibrio parahaemolyticus *(Vp*-FlhA) and SctV from Yersinia enterocolitica (*Ye*-SctV) for further studies, as both could be produced in large quantities at high purity (Fig 1b) using the gentle detergent LMNG. When the purified proteins were subjected to gel filtration, both produced high molecular weight complexes (Fig 1c).

**Figure 1.**
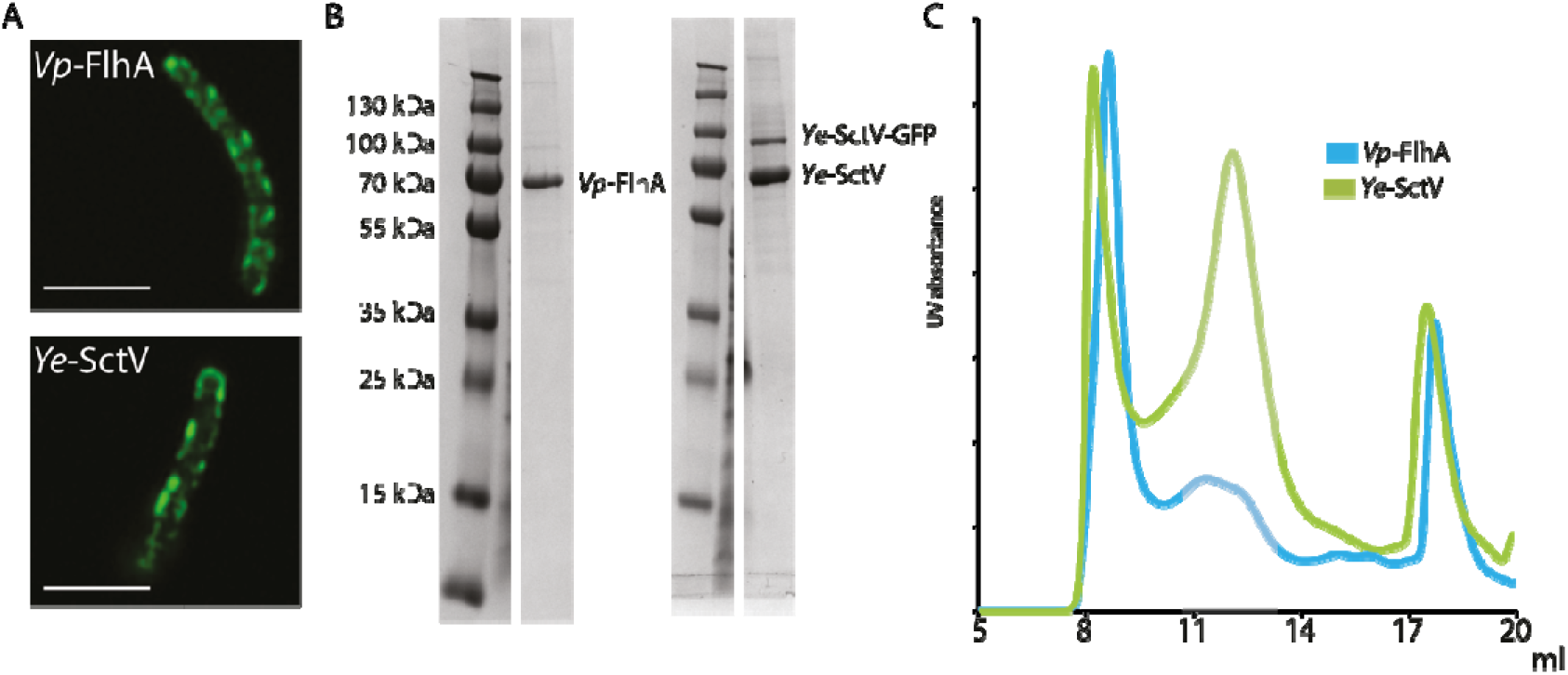
FlhA from Vibrio parahaemolyticus and SctV from Yersinia enterocolitica assemble into a large complex in the membrane of E. coli. **a.** Fluorescence images of *E. coli* cells expressing GFP tagged *Vp*-FlhA or *Ye*-SctV in the cell envelope. Scale bar 5 μm. **b.** SDS-PAGE of purified *Vp*-FlhA and *Ye*-SctV following cleavage of GFP. **c.** Gel filtration traces of purified *Vp*-FlhA and *Ye*-SctV. The grey shaded area indicates fractions collected to make cryoEM grids.

We imaged the purified *Vp*-FlhA and *Ye*-SctV complexes using cryo-EM and analysed them using single particle analysis in Relion (29). Particle classification revealed that *Ye*-SctV had formed a dimer of nonamers and we imposed D9 symmetry for 3D refinement, resulting in a 3.7 Å reconstruction of the cytoplasmic domain double nonamer (Fig 2a). No high-resolution features of the transmembrane domain could be discerned. The *Ye*-SctV cytoplasmic domain 18-mer is made up of two nonamers dimerising via the cytoplasm facing side of the cytoplasmic domain. This assembly is not consistent with the known localisation of SctV in the injectisome (30) and is therefore considered to be an artefact of the removing the complex from the constraint of the membrane.

**Figure 2.**
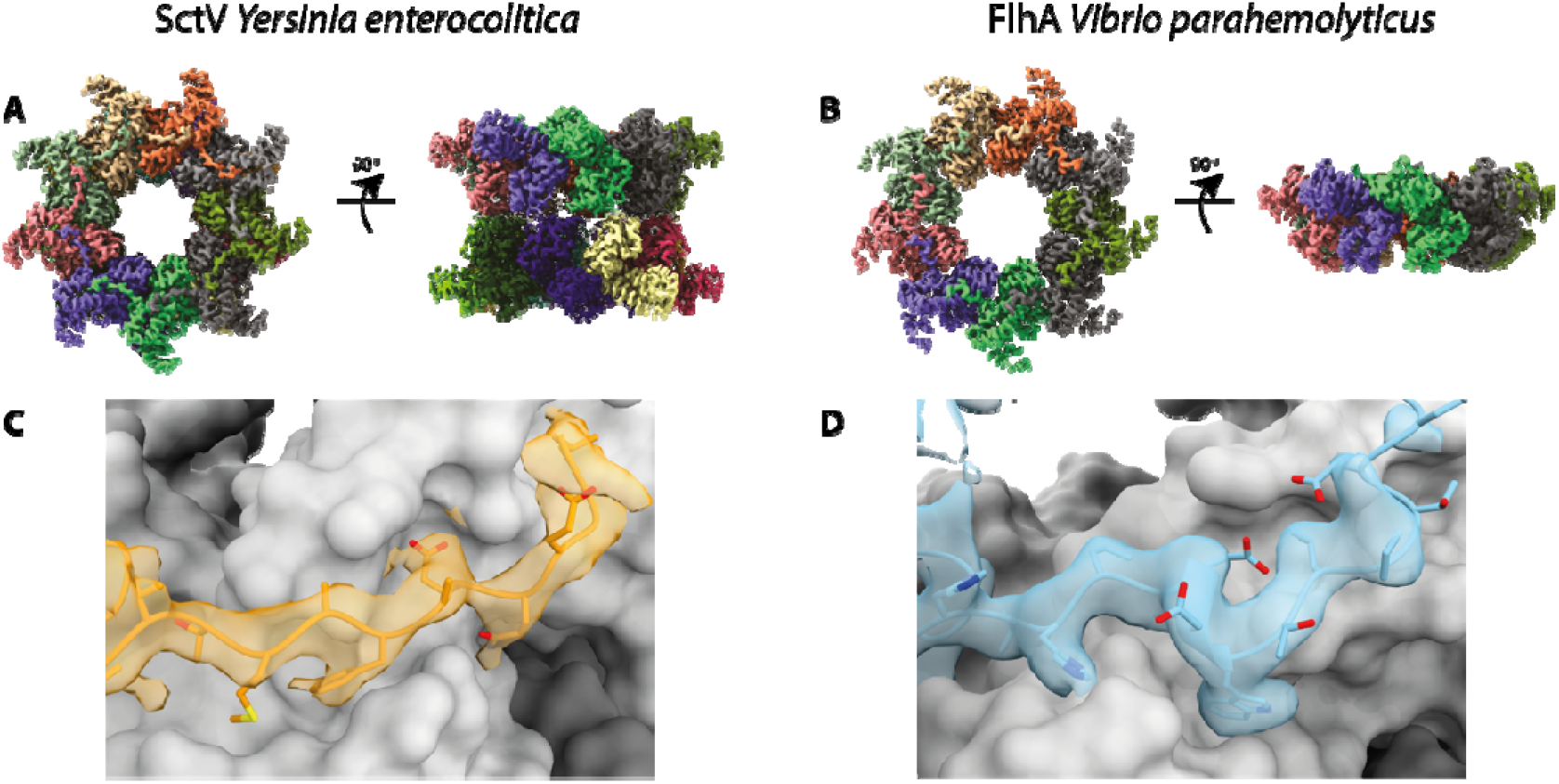
Cryo-EM volumes of the cytoplasmic domain of SctV and FlhA. **a.** 3D reconstruction of *Ye*-SctV at a resolution of 3.7 Å using D9 symmetry. **b.** 3D reconstruction of *Vp*-FlhA at a resolution of 3.8 Å using C9 symmetry. **c.** Model of the *Ye*-SctV linker region in the cryo-EM volume (orange) in the context of the adjacent protomer (grey). **d.** Model of the *Vp*-FlhA linker region in the cryo-EM volume (blue) in the context of the adjacent protomer (grey).

In contrast, *Vp*-FlhA was found to form the predicted nonameric complex and we imposed C9 symmetry for particle refinement, resulting in a 3.8 Å volume of the cytoplasmic domain but little detail in the transmembrane domain (Fig 2b). Both reconstructions were of sufficient quality to allow us to build a model of the structure of the cytoplasmic domain.

A striking feature of both structures is the N-terminal density of the cytoplasmic domain, formed by FlhA_L_/SctV_L_, which reaches out to bind to the adjacent protomer (Fig 2c), demonstrating this well studied interaction for the first time in the context of the full-length protein in both flagellar and injectisome T3SS. Biochemical studies have indicated that the interaction between FlhA_L_ and the neighbouring subunit is important for assembly of the nonameric ring (17), however, the interaction has so far only been observed in crystal structures of monomeric protein (20, 21) and SctV_L_ is not ordered in the crystal structure of the assembled *Sf*-SctV nonamer (11). In both of our structures a hydrophobic residue, Trp350 in *Vp*-FlhA and Phe367 in *Ye*-SctV, sticks into a hydrophobic groove on the neighbouring subunit. This residue in FlhA has previously been shown to be involved in ring formation of the cytoplasmic domain of FlhA (17).

The overall dimensions of the nonameric rings are similar to that of the crystal structure of *Sf*-SctV (Fig 3a), but the individual subunits are in a different state. FlhA/SctV is known to be able to adopt at least two conformations, open and closed, and FlhA_L_/SctV_L_ is thought to bind to the membrane proximal face of the neighbouring subunit only in the open state (19). Comparison of our structures with FlhA from *S.* Typhimurium, *St*-FlhA, in the open state and *Sf*-SctV, which is in the closed state shows that both of our complexes are in the open state (Fig 3b), with *Ye*-SctV being the first example of an SctV homologue in the open state. The demonstration of the open state opens up the possibility that SctV binds to substrate-chaperone complexes in an analogous manner to that described for FlhA (18), in which chaperones bind between Sub-Domain(SD)4 and SD2 and are sterically unable to access the binding site in the closed state.

**Figure 3.**
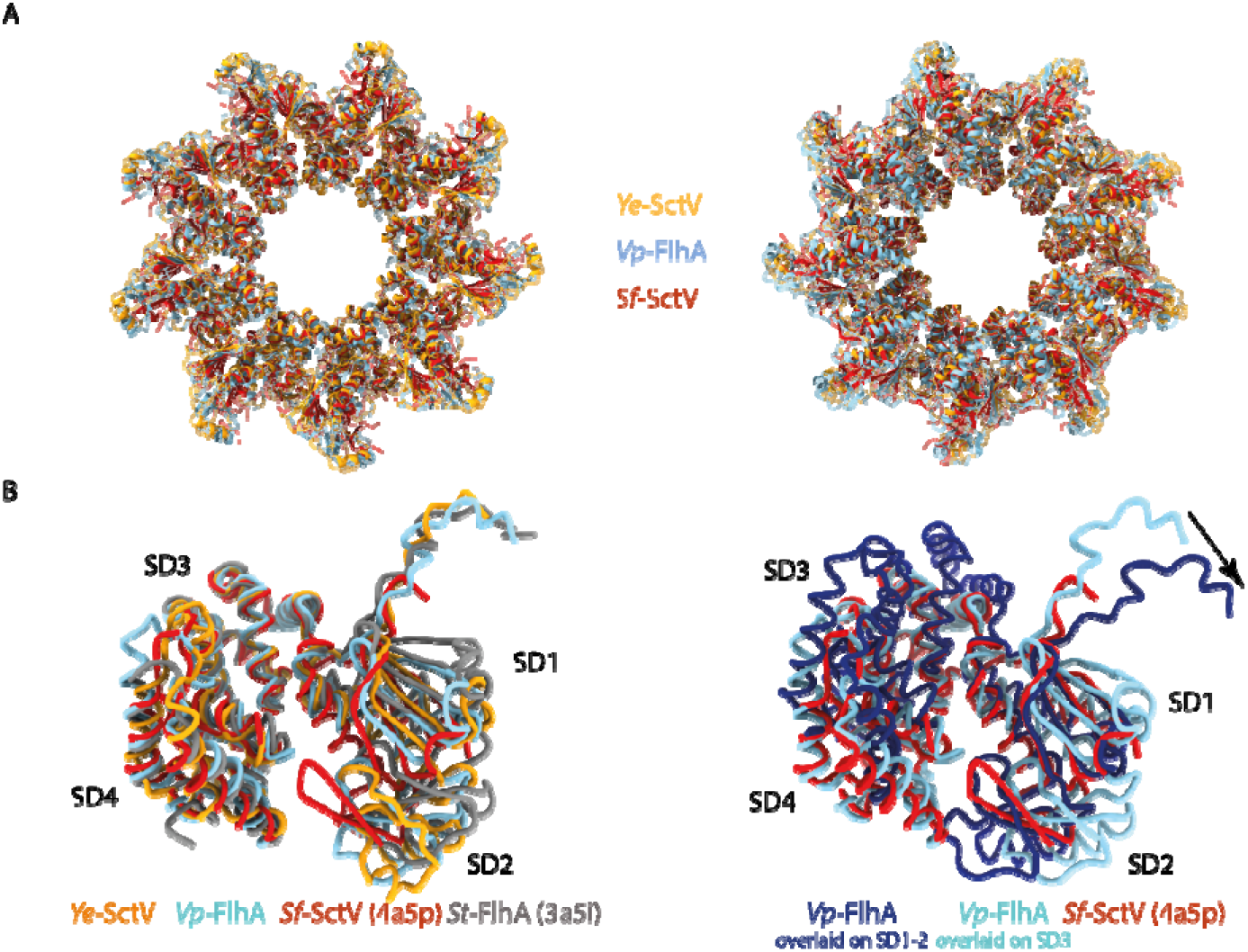
The cryo-EM structures of Ye-SctV and Vp-FlhA are in the open state. **a.** Overlay of the nonameric rings of the cytoplasmic domains of Ye-SctV and Vp-FlhA with the crystal structure of the *Sf*-SctV nonameric ring (PDB: 4A5P). **b.** Overlay of a single subunit of *Ye*-SctV and *Vp*-FlhA with *St*-FlhA in the open state (grey, PDB: 3A5I) and *Sf*-SctV in the closed state (red, PDB: 4A5P). **c.** Overlay of *Vp*-FlhA (light blue) on the SD3 domain of *Sf*-SctV (red) and overlay (dark blue) on the SD1 and SD2 domains of *Sf*-SctV illustrating the movement of the linker (arrow) as the protein changes from the open to the closed state.

A conformational change from open to closed state would be expected to lead to further changes throughout the protein. We simulated the movement of FlhA_L_ by overlaying one subunit of *Vp*-FlhA onto the SD3 or SD2 subdomain of the closed *Sf*-SctV structure (Fig 3c). This demonstrates that adoption of the closed state in one subunit would affect the neighbouring subunit, potentially propagating the conformational change in a wave around the ring. An additional effect on the transmembrane domain is possible.

## Discussion

Full-length FlhA/SctV forms a very fragile, detergent sensitive membrane protein complex. Here, we have identified more stable FlhA/SctV sequences and purification conditions that have allowed preparation of the assembled ring complex for structural studies. We have determined nonameric structures of the cytoplasmic domain of FlhA/SctV in the context of the full-length protein and in the open state. Multiple studies have previously implicated the FlhA linker region, FlhA_L_, in promoting complex formation. We provide the first structural view of this interaction in the assembled state in both FlhA and SctV, showing that SctV_L_ is the functional equivalent of FlhA_L_ in the stabilisation of the nonameric assembly.

It has been proposed that the closed state of the FlhA/SctV cytoplasmic domain interacts with early secretion substrates, while later substrates bind to the open state (19) which is induced by completion of the first step of filament assembly. Both of our structures are in the open state. Our structures are not affected by crystallisation artefacts and are in the context of the full-length protein, suggesting that the open state is the default state in the absence of other factors. Our structures also suggest that at least in the open state the residues known as FlhA_L_/SctV_L_, which have previously been described as the linker between cytoplasmic and membrane domains, do not form part of this linker but are part of the cytoplasmic domain. Instead, the residues N-terminal to FlhA_L_/SctV_L_ must be responsible for linking the membrane domain to the cytoplasmic domain.

Unfortunately, we were not able to determine the structure of the membrane domain of either complex, possibly due to damage to the fragile membrane domain at the air-water interface during cryo-EM sample preparation (31). However, two features are clear in 2D averages of the side view of both complexes. The cytoplasmic and membrane domains are at a set distance from each other (Fig 4b) and this distance matches that observed in tomographic reconstructions of the T3SS (Fig 4a). This suggests that the linker between the domains is not flexible, which would be inconsistent with a previously suggested model of secretion according to which the cytoplasmic domain is tethered to the membrane by a flexible linker and moves towards the membrane domain and back into the cytoplasm in a cyclical fashion (23).

**Figure 4.**
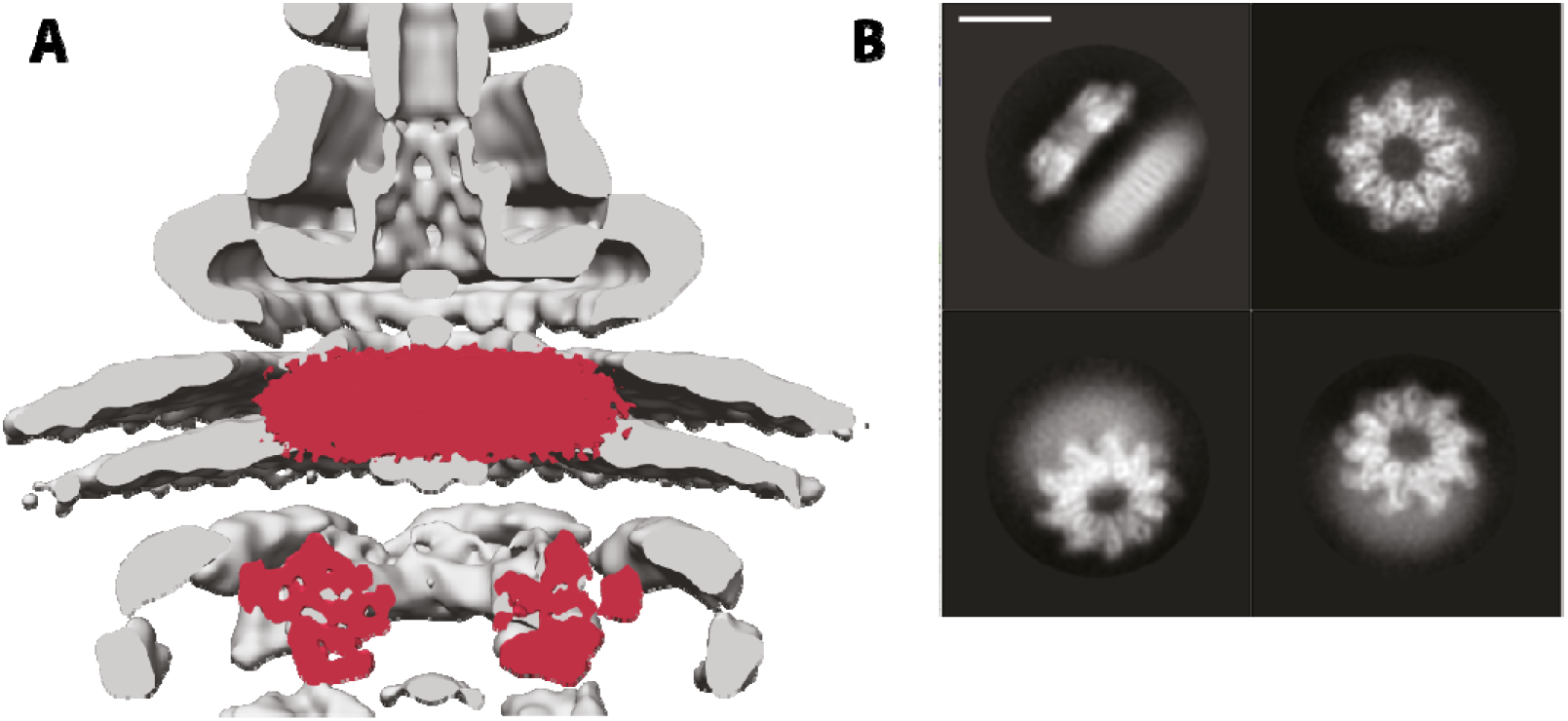
Position of the membrane domain of FlhA in the T3SS nanomachine. **a.** The volume of *Vp*-FlhA is shown in red overlaid onto the tomographic reconstruction of the injectisome (EMD-8544). **b.** Selected 2D class averages of *Vp*-FlhA show the high level of detail in the cytoplasmic domain and the lower information content in the presumed transmembrane, micelle-embedded, portion. Scale bar is 100A.

While this manuscript was in preparation, a cryo-EM structure of the nonameric cytoplasmic domain of *E. coli* SctV in the closed state (32) and crystal structures of the nonameric cytoplasmic domain of *Chlamydia pneumoniae* SctV alone and in complex with the ATPase stalk SctO were published (33). While molecular dynamics simulations suggested that *E. coli* SctV does not easily switch between open and closed state, our SctV and the *C. pneumoniae* SctV structure show that SctV can exist in the open state. The closed state may be favoured in some crystallisation conditions. Interestingly, the *C. pneumoniae* SctV conformation is altered by the binding of the SctO stalk protein, suggesting a mechanism for modifying the substrate-chaperone binding sites during the secretion cycle from SctV.

The methods developed here and the identification of FlhA and SctV homologues that can assemble in the membrane in the absence of other T3SS components may be the basis of future studies of the FlhA/SctV transmembrane domain or interactions of the cytoplasmic domain with chaperones and substrates in the context of the assembled complex.

## Methods

### Protein purification

FlhA and SctV were produced as GFP fusion proteins in *E. coli* BL21 by expressing them from a pt12 plasmid (5). Cells were grown in terrific broth supplemented with rhamnose monohydrate (0.1%) and harvested after overnight growth. Cells were resuspended in TBS (100 mM Tris, 150 mM NaCl, 1 mM EDTA) and lysed using an Emulsiflex homogeniser (Avestin). Membranes were purified by ultracentrifugation of the clarified lysate at 235,000g and solubilised in 1% LMNG using 0.1 grams of detergent per gram of membrane pellet. After gentle stirring for one hour aggregates were removed by centrifugation at 75,600g and the solubilised membrane proteins were applied to a StrepTrap column (GE healthcare) which was then washed in TBS supplemented with 0.01% LMNG. Pure protein was eluted in TBS containing 0.01% LMNG and 10 mM desthiobiotin. GFP was cleaved off by incubating with tev protease overnight. Next, the protein was concentrated and further purified by gel filtration using a superpose 6 increase (GE healthcare) in TBS containing 0.01% LMNG. The peak fraction, eluting close to 12 ml, was collected.

### Cryo-EM analysis

3 μl of purified protein at a concentration between 0.8 and 1 mg/ml was deposited on a glow-discharged Quantifoil grid (2/2, 300 mesh for FlhA, 1.2/1.3, 300 mesh for SctV). Grids were blotted and plunged into liquid ethane using a Vitrobot Mark IV (FEI). The samples were imaged using a Titan Krios (FEI) equipped with a K2 detector (Gatan). Relion’s implementation of MotionCor2 (34) was used for motion correction and SIMPLE’s implementation of CTFFIND4 was used for CTF estimation (35). Particles were picked using SIMPLE and classified in Relion3.0 (29). Atomic models were built in Coot (36) and refined using phenix (37).

### Fluorescence imaging

8 μl of an overnight culture of *E coli* BL21 expressing FlhA-GFP or SctV-GFP were applied to a glass slide and imaged using a Zeiss 880 inverted microscope equipped with a plan-apochromat 63x/1.4 NA objective and an Airyscan detector. GFP fluorescence was excited using a laser (488 nm).

## Acknowledgements

We thank Jerry Cao for ably assisting experiments during his Part II project work, the staff of the Central Oxford Structural Microscopy and Imaging Centre Errin Johnson and Adam Costin, Alan Wainman of the Dunn School Light Microscopy Facility and all members of the Lea group for assistance. The work was funded by grants from the Wellcome Trust 109136, 100298, 219477, the Medical Research Council (UK) S021264 and the Wolfson Foundation WL160052. Work at the CSB was funded by core funding from the CCR.

## Data Availability

Coordinates and associated volumes and metadata have been deposited in the PDB and EMDB respectively. SctV EMD-11820 / PDB 7ALW, FlhA EMD-11827 / PDB 7AMY.

---

**Table 1.**
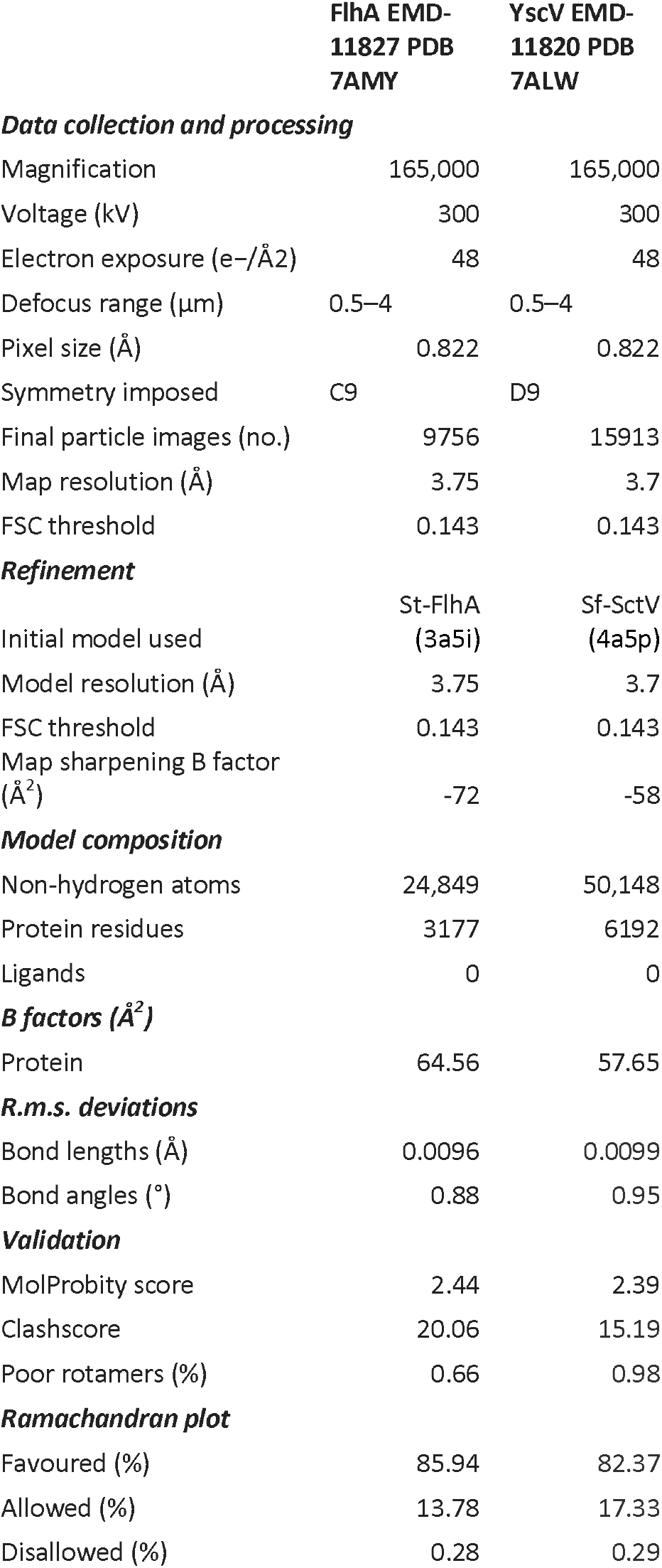
CryoEM data collection, processing and model statistics

|  | FlhA EMD-11827 PDB<br>7AMY | YscV EMD-11820 PDB<br>7ALW |
| --- | --- | --- |
| **Data collection and processing** |  |  |
| Magnification | 165,000 | 165,000 |
| Voltage (kV) | 300 | 300 |
| Electron exposure (e-/Å<sup>2</sup>) | 48 | 48 |
| Defocus range (μm) | 0.5–4 | 0.5–4 |
| Pixel size (Å) | 0.822 | 0.822 |
| Symmetry imposed | C9 | D9 |
| Final particle images (no.) | 9756 | 15913 |
| Map resolution (Å) | 3.75 | 3.7 |
| FSC threshold | 0.143 | 0.143 |
| **Refinement** |  |  |
| Initial model used | St-FlhA<br>(3a5i) | Sf-SctV<br>(4a5p) |
| Model resolution (Å) | 3.75 | 3.7 |
| FSC threshold | 0.143 | 0.143 |
| Map sharpening B factor<br>(Å<sup>2</sup>) | -72 | -58 |
| **Model composition** |  |  |
| Non-hydrogen atoms | 24,849 | 50,148 |
| Protein residues | 3177 | 6192 |
| Ligands | 0 | 0 |
| **B factors (Å<sup>2</sup>)** |  |  |
| Protein | 64.56 | 57.65 |
| **R.m.s. deviations** |  |  |
| Bond lengths (Å) | 0.0096 | 0.0099 |
| Bond angles (°) | 0.88 | 0.95 |
| **Validation** |  |  |
| MolProbity score | 2.44 | 2.39 |
| Clashscore | 20.06 | 15.19 |
| Poor rotamers (%) | 0.66 | 0.98 |
| **Ramachandran plot** |  |  |
| Favoured (%) | 85.94 | 82.37 |
| Allowed (%) | 13.78 | 17.33 |
| Disallowed (%) | 0.28 | 0.29 |

## Notes

### Competing Interest Statement

The authors have declared no competing interest.

